# A divergent *Articulavirus* in an Australian gecko identified using meta-transcriptomics and protein structure comparisons

**DOI:** 10.1101/2020.05.21.109603

**Authors:** Ayda Susana Ortiz-Baez, John-Sebastian Eden, Craig Moritz, Edward C. Holmes

**Affiliations:** Marie Bashir Institute for Infectious Diseases and Biosecurity, School of Life and Environmental Sciences and School of Medical Sciences, The University of Sydney, Sydney, New South Wales 2006, Australia; Centre for Virus Research, Westmead Institute for Medical Research, Westmead, NSW 2145, Australia; Research School of Biology & Centre for Biodiversity Analysis, The Australian National University, Acton, ACT 6201, Australia

**Keywords:** virus discovery, protein structure, meta-transcriptomics, *Tilapia tilapinevirus*, *Articulavirales*, *Amnoonviridae*, RNA virus, gecko

## Abstract

The discovery of highly divergent RNA viruses is compromised by their limited sequence similarity to known viruses. Evolutionary information obtained from protein structural modelling offers a powerful approach to detect distantly related viruses based on the conservation of tertiary structures in key proteins such as the viral RNA-dependent RNA polymerase (RdRp). We utilised a template-based approach for protein structure prediction from amino acid sequences to identify distant evolutionary relationships among viruses detected in meta-transcriptomic sequencing data from Australian wildlife. The best predicted protein structural model was compared with the results of similarity searches against protein databases based on amino acid sequence data. Using this combination of meta-transcriptomics and protein structure prediction we identified the RdRp (PB1) gene segment of a divergent negative-sense RNA virus in a native Australian gecko (*Geyra lauta*) that was confirmed by PCR and Sanger sequencing. Phylogenetic analysis identified the *Gecko articulavirus* (GECV) as a newly described genus within the family *Amnoonviridae*, order *Articulavirales*, that is most closely related to the fish virus *Tilapia tilapinevirus* (TiLV). These findings provide important insights into the evolution of negative-sense RNA viruses and structural conservation of the viral replicase among members of the order *Articulavirales*.

## Introduction

The development of next-generation sequencing technologies (NGS), including total RNA sequencing (meta-transcriptomics), has revolutionized studies of virome diversity and evolution [1–3]. Despite this, the discovery of highly divergent viruses remains challenging because of the often limited (or no) primary sequence similarity between putative novel viruses and those for which genome sequences are already available [4–6]. For example, it is possible that the small number of families of RNA viruses found in bacteria, as well as their effective absence in archaeabacteria, in reality reflects the difficulties in detecting highly divergent sequences rather than their true absence from these taxa [3].

The conservation of protein structures in evolution and the limited number of proteins folds (fold space) in nature form the basis of template-based protein structure prediction [7], providing a powerful way to reveal the origins and evolutionary history of viruses [8,9]. Indeed, the utility of protein structural similarity in revealing key aspects of virus evolution is well known [9,10]. For instance, double-strand (ds) DNA viruses including the thermophilic archaeal virus STIV, enterobacteria phage PRD1, and human adenovirus exhibit conserved viral capsids, suggesting a deep common ancestry [11]. Thus, protein structure prediction utilising comparisons to solved protein structures can assist in the identification of potentially novel viruses [7,12]. Herein, we use this method as an alternative approach to virus discovery.

There is a growing availability of three-dimensional structural data in curated databases such as the Protein Data Bank (PDB), with approximately 11,000 viral protein solved structures that can be used in comparative studies. Importantly, these include structures of the RNA-dependent RNA polymerase (RdRp) that exhibits the highest level of sequence similarity among RNA viruses, including a number of key conserved motifs, and hence is expected to contain relatively well conserved protein structures. Exploiting such structural features in combination with metagenomic data will undoubtedly improve our ability to detect divergent viruses in nature, particularly in combination with wildlife surveillance [2,4,13].

The International Committee on Taxonomy of Viruses (ICTV) recently introduced the *Amnoonviridae* as a newly recognized family of negative-strand RNA viruses present in fish (ICTV Master Species List 2018b.v2). Together with the *Orthomyxoviridae*, the *Amnoonviridae* are classified in the order *Articulavirales*, describing a set of negative-sense RNA viruses with segmented genomes. While the *Orthomyxoviridae* includes seven genera, four of these comprise influenza viruses (FLUV), and to date the family *Amnoonviridae* comprises a single genus – *Tilapinevirus* – which in turn includes only a single species - *Tilapia tilapinevirus* or Tilapia Lake virus (TiLV).

TiLV was originally identified in farmed tilapine populations (*Oreochromis niloticus*) in Israel and Ecuador [14]. The virus has now been described in wild and hybrid tilapia across several countries in the Americas, Africa, Asia, and Southeast Asia [15–17]. TiLV has been associated with high morbidity and mortality in infected animals. Pathological manifestations include syncytial hepatitis, skin erosion and encephalitis [15,18]. TiLV was initially classified as a putative orthomyxo-like virus based on weak sequence resemblance (~17% amino acid identity) in the PB1 segment that contains the RdRp, as well as the presence of conserved 5′ and 3′ termini [14]. While both the *Orthomyxoviridae* and *Amnoonviridae* have negative-sense, segmented genomes, the genomic organization of the *Amnoonviridae* comprises 10 instead of 7-8 segments [14,18,19], and their genomes are shorter (~10 kb) than those of the *Orthomyxoviridae* (~12-15 kb). To date, however, only the RdRp (encoded by a 1641 bp PB1 sequence) has been reliably defined, and most segments carry proteins of unknown function. Importantly, comparisons of TiLV RdRp with sequences from members of the *Orthomyxoviridae* revealed the presence of four conserved amino acid motifs (I-IV) of size 4-9 amino acid residues each [14] that effectively comprise a “molecular fingerprint” for the order.

Unlike other members of the *Articulavirales* [20], TiLV appears to have a limited host range and has been only documented in tilapia (*O. niloticus*, *O*. sp.) and hybrid tilapia (*O. niloticus* x *O. aureus*). Herein, we report the discovery of a divergent virus from an Australian gecko (*Geyra lauta*) using a combination of meta-transcriptomic and structure-based approaches, and employ a phylogenetic approach to reveal its relationship to TiLV. Our work suggests that this Gecko virus likely represents a novel genus within the *Amnoonviridae*.

## Materials and Methods

### Sample collection

A total of seven individuals corresponding to the reptile species *Carlia amax, Carlia gracilis, Carlia munda, Gehyra lauta, Gehyra nana, Heteronotia binoei*, and *Heteronotia planiceps* were collected alive in 2013 from Queensland, Australia. Specimens were identified by mtDNA typing and/or morphological data. Livers were harvested and stored in RNAlater at −80°C before downstream processing. All sampling was conducted in accordance with animal ethics approval (#A2012/14) from the Australian National University and collection permits from the Parks and Wildlife Commission of the Northern Territory (#45090), the Australian Government (#AU-COM2013-192), and the Department of Environment and Conservation (#SF009270).

### Sampling processing and sequencing

RNA extraction was performed using the RNeasy Plus minikit (Qiagen) following manufacturer’s instructions. Each of the seven livers were extracted individually and then pooled in equal amounts. For RNA sequencing, ribosomal RNA (rRNA) was depleted using the RiboZero (epidemiology) depletion kit and libraries were prepared with the TruSeq stranded RNA library prep kit before sequencing on an Illumina HiSeq 2500 platform (100 bp paired end reads). Library preparation and sequencing was performed by the Australian Genome Research Facility (AGRF), generating a total of 22,394,787 paired end reads for the pooled liver RNA library.

### De novo assembly and sequence annotation

Raw Illumina reads were trimmed of sequencing adapters and low-quality bases with Trimmomatic v0.38 [21]. The trimmed reads were then *de novo* assembled into contigs (transcripts) using Trinity v2.8.6 [22]. Contig abundance was estimated with RSEM [23] and shown as the numbers of transcripts per million (TPM). For sequence annotation, contigs were compared against the NCBI nucleotide (nt) and non-redundant (nr) protein databases (nr) using BLASTn [24] and DIAMOND [25], respectively.

### Protein structure prediction for virus detection

To further screen the meta-transcriptomic data, all the assembled sequences below the assigned threshold (e-value ≥ 10^−5^) were assigned as “orphan” contigs (n= 293,586). These were then analysed using a protein structure-informed approach. Specifically, orphan contigs were translated into all six open reading frames (ORFs) using the getorf program [26] to identify continuous ORFs of at least 1000nt in length between two stop codons (n=57). To detect distant sequence homologies and predict viral protein structures, this subset of translated ORFs were then analysed using a template-based modelling approach as implemented in Phyre2 (http://www.sbg.bio.ic.ac.uk/phyre2) [27]. In brief, target proteins were compared against proteins of known structure via homology modelling and fold recognition, followed by loop modelling and sidechain fitting [27]. Confident matches (confidence >90%) to known viral structures were selected for downstream analyses. Annotations from the predicted model were used as preliminary data for tentative taxonomic assignment and protein classification.

### Annotation of the newly discovered virus

To further corroborate the viral origin of the predicted protein structure and gain insights into its taxonomic classification, we conducted parallel comparisons using DIAMOND [25] against the GenBank non-redundant (nr) database (https://www.ncbi.nlm.nih.gov/) and the HMMER web server (http://www.ebi.ac.uk/Tools/hmmer) against the following profile databases: (i) reference proteomes (https://proteininformationresource.org/rps/), (ii) Uniprot (https://www.uniprot.org/) and (iii) Pfam (https://pfam.xfam.org/). In addition, conserved domains were annotated using the Conserved Domain Database (CDD) and the CD-search tool (http://www.ncbi.nlm.nih.gov/Structure/cdd/cdd.shtml). To detect additional contigs and better characterize the entire genome of the novel virus, we aligned the DNA contigs against custom databases using DIAMOND [25], including (i) a reference RdRp sequences from the order *Articulavirales*, and (ii) reference sequences corresponding to all the segments of TiLV (Table S1). Given the divergent nature of the viruses, we considered all hits with E-value >10^−4^.

### Phylogenetic analysis

The predicted contig encoding the RdRp of the newly discovered virus was aligned with reference protein sequences of the order *Articulavirales* (Table S2). A multiple amino acid sequence alignment was performed using the E-INS-i algorithm as implemented in the MAFFT v7.450 program [28]. Selection of the best-fit model of amino acid substitution was carried out using the Akaike Information criterion (AIC) and the Bayesian Information Criterion (BIC) with the standard model selection option (-m TEST) in IQ-TREE [29]. Phylogenetic analysis of these data was then performed using the Maximum Likelihood (ML) method available in IQ-TREE, with node support estimated with the ultra-fast bootstrap (UFBoot) approximation (1000 replicates) and the Shimodaira-Hasegawa approximate Likelihood ratio test (SH-aLRT). Sequencing reads are available at the NCBI Sequence Read Archive (SRA) under the Bioproject PRJNA626677 (BioSample: SAMN14647831; Sample name: VERT7; SRA: SRS6507258). The assembled sequence for GECV was deposited in GenBank under the accession number MT386081.

### PCR validation

To validate the presence of the novel gecko amnoonvirus, and to identify the putative host species, we screened the individual liver RNA using RT-PCR. Briefly, cDNA was prepared using Superscript IV VILO master mix and RT-PCR was performed with the Platinum SuperFi Green PCR master mix and two primers sets targeting the gecko RdRp contig – F2V7 and F3V7 (Table S3). The resultant RT-PCR products were analysed by agarose gel electrophoresis and validated by Sanger sequencing.

## Results

### Virus discovery using meta-transcriptomics and protein structural features

We used a meta-transcriptomic approach to screen a single pooled library containing liver RNA of seven Australian native reptile species (*Gehyra lauta, Carlia amax, Heteronotia binoei, Gehyra nana, Carlia gracilis, Carlia munda*, and *Heteronotia planiceps*; see Methods). We focused on the *de novo* assembled contigs that had no significant hits using initial searches against the NCBI nucleotide and non-redundant databases. Accordingly, of 293,586 orphan contigs, 57 contained translatable ORFs of more than 1000 nt in length, and because we hypothesized that some may correspond to undetected virus sequences, we interrogated them using a protein structure prediction approach with template-based modelling (TBM) in Phyre2 [27]. From the 57 queried contigs, we obtained a 3D model of a 407 amino acid (1227 bp) contig with a high confidence hit (98.3%) to the RdRp catalytic subunit of a bat influenza A virus (family *Orthomyxoviridae*) (Table 1, Figure 1a-b). The confidence level obtained is indicative of high probability of modelling success between putative homologs. In addition, the alignment coverage between our query and the viral template corresponded to 52% (213 residues) of the query sequence, while the proportion of identical amino acids (i.e. sequence identity) was 19% (Table 1).

**Table 1.** Summary of analyses and parameters used for the detection of GECV.

| Analysis/database | Parameter (unit) | Value / Hit (e-value) |
| --- | --- | --- |
| **Trinity *de novo* assembly** | Length (nt) | 1227 |
|  | Predicted ORF length (aa) | 407 |
|  | Coverage (# of reads) | 35 |
|  | Abundance (TPM <sup>1</sup>) | 1.10 |
| **Phyre2/PDB** | PDB molecule | RdRp catalytic subunit |
|  | PDB title | Bat influenza a polymerase with bound vRNA promoter |
|  | PDB identifier | 4WSB |
|  | Resolution | 2.65 |
|  | Confidence (%) | 98.3 |
|  | Coverage (%) | 52 |
|  | Identity (%) | 19 |
| **DIAMOND/nr** | Match | Hypothetical protein (Tilapia lake virus), segment 1 |
|  | Similarity (%) | 29 |
|  | E-value | 1.30E-07 |
| **DIAMOND/custom db** | Match | Hypothetical protein (Tilapia lake virus), segment 1 |
|  | Similarity (%) | 29 |
|  | E-value | 2.4E-14 |
| **HMMER/references proteomes** | Taxonomy | Tilapia lake virus (3.9e-11) |
|  | Domain architecture | Flu_PB1 |
| **HMMER/UniProt** | Taxonomy | Tilapia lake virus (1.4e-10) |
|  | Domain architecture | Flu_PB1 |
| **HMMER/SwissProt** | Taxonomy | Infectious salmon anaemia virus RDRP_ISAV8, segment 2 (5.2e-3) |
|  | Domain architecture | Flu_PB1 |
| **Pfam** | Family | Flu_PB1 (1.8e-2) |
|  | Description | Influenza RNA-dependent RNA polymerase subunit PB1 |
| **CDD/CDDv3.17** | Domain hit | Flu_PB1 super family (6.43e-05) |

<sup>1</sup>TPM: transcripts per million.

**Figure 1.**
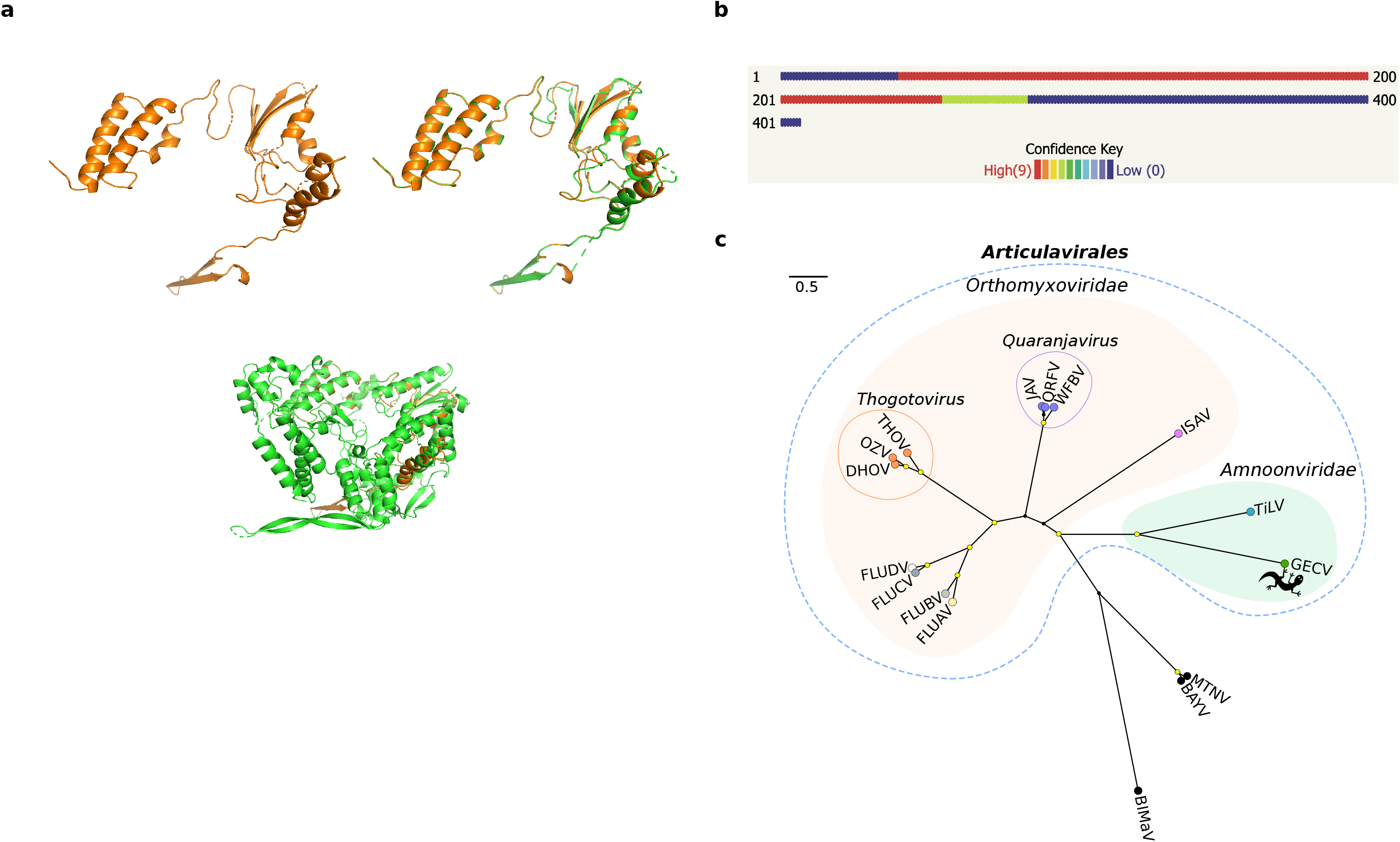
Protein structure prediction and phylogenetic relationships of GECV. (**a**) 3D model prediction of the RdRp subunit PB1 of GECV (top left). Protein structure superposition in the aligned region between the predicted model for GECV and the RdRp (PB1 gene) of influenza A virus (FLUAV) (top right). Protein structure superposition of the predicted model for GECV and the entire RdRp subunit of FLUAV (bottom). The protein structure predicted for GECV is displayed in orange and that of FLUAV in green. (**b**) Confidence summary of residues modelled. (**c**) Maximum likelihood tree depicting the phylogenetic relationships between GECV and TiLV within the family *Amnoonviridae*, order *Articulavirales*. Families are indicated with colored filled bubbles. Tip labels are colored according to genus. Genera comprising multiple species are indicated with unfilled bubbles. Support values >= 95% UFBoot and 80% SH-aLRT are displayed with yellow-circle shapes at nodes. *Alphainfluenzavirus* (FLUBA); *Betainfluenzavirus* (FLUBV); *Deltainfluenzavirus* (FLUDV); *Gammainfluenzavirus* (FLUCV); *Dhori thogotovirus* (DHOV); Oz virus (OZV); *Thogoto thogotovirus* (THOV); *Quaranfil quaranjavirus* (QRFV); *Wellfleet Bay virus* (WFBV); *Johnston Atoll quaranjavirus* (JAV); *Salmon isavirus* (ISAV); *Tilapia tilapinevirus* (TiLV); *Gecko articulavirus* (GECV); *Blueberry mosaic associated virus* (BIMaV); *Montano orthohantavirus* (MTNV); *Bayou orthohantavirus* (BAYV).

To corroborate these findings, the structural results were compared with those obtained from other analyses based on primary sequence similarity searches against public databases (see Methods) (Table 1). This revealed matches to the RdRp subunit (PB1 gene segment) of different members of the order *Articulavirales*, including the Influenza virus (FLUAV), TiLV, and Infectious salmon anaemia virus (ISAV). Comparisons of the assembled contigs against a custom database containing only members of the *Articulavirales* were then performed to improve sequence alignments. Accordingly, the best hit matches were obtained to TiLV (e-values <10^−15^) (Table 1). To identify additional viral segments, the assembled contigs were aligned to the ten segments of TiLV using DIAMOND. A total of 87 contigs were scored through the entire genome, although we did not recover any significant hit for segments 2-10 likely because they are so divergent in sequence (Table S1).

### Sequence alignment and phylogenetic relationships

We tentatively name the new virus identified here as Gecko articulavirus (GECV). Multiple sequence alignment of the RdRp between GECV and other members the order *Articulavirales* identified a number of well conserved amino acid motifs (I-IV) ranging in length from 5-11 amino acids in length (Figure 2). Phylogenetic analysis of the aligned RdRp region revealed that GECV falls within the order *Articulavirales* and, along with TiLV (family *Amnoonviridae*), comprises a distinct monophyletic group. The close relationship between GECV and TiLV was supported by high UFBoot/SH-aLRT values (99%/99%) (Figure 1c). Likewise, estimates of the amino acid identity in the RdRp showed a closer (but still distant) sequence similarity (15.35%) with TiLV than other members of the order *Articulavirales* (Table 2).

**Table 2.**
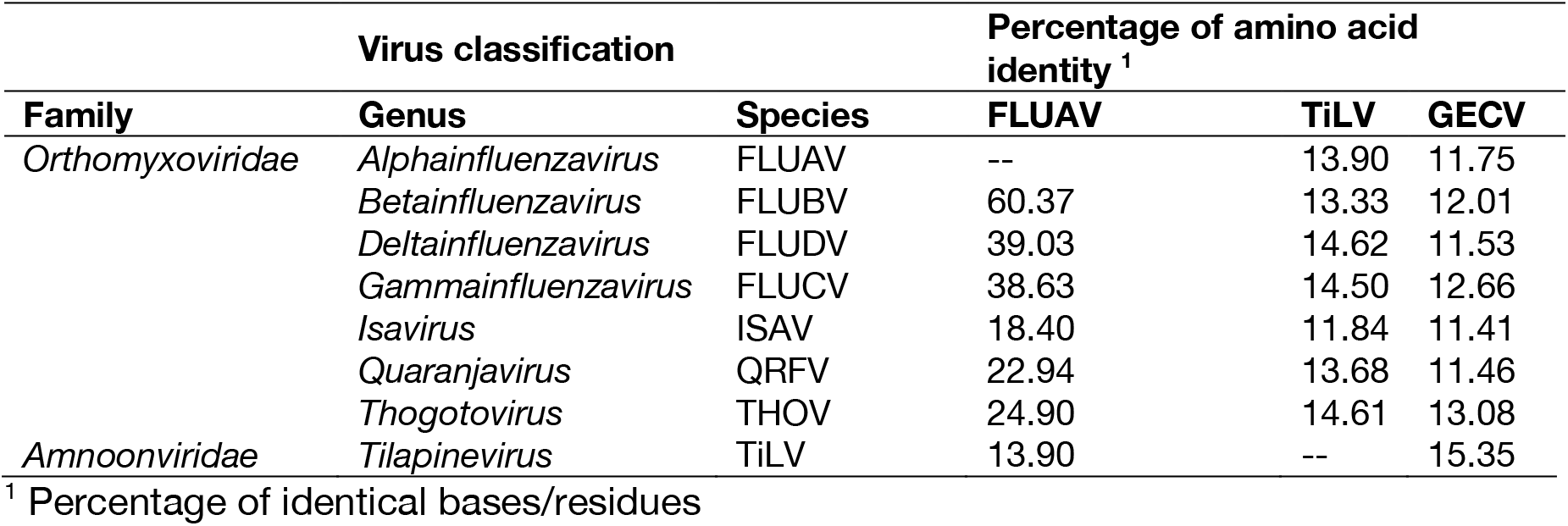
Percentage of identical residues among members of the order *Articulavirales* and GECV.

**Figure 2.**
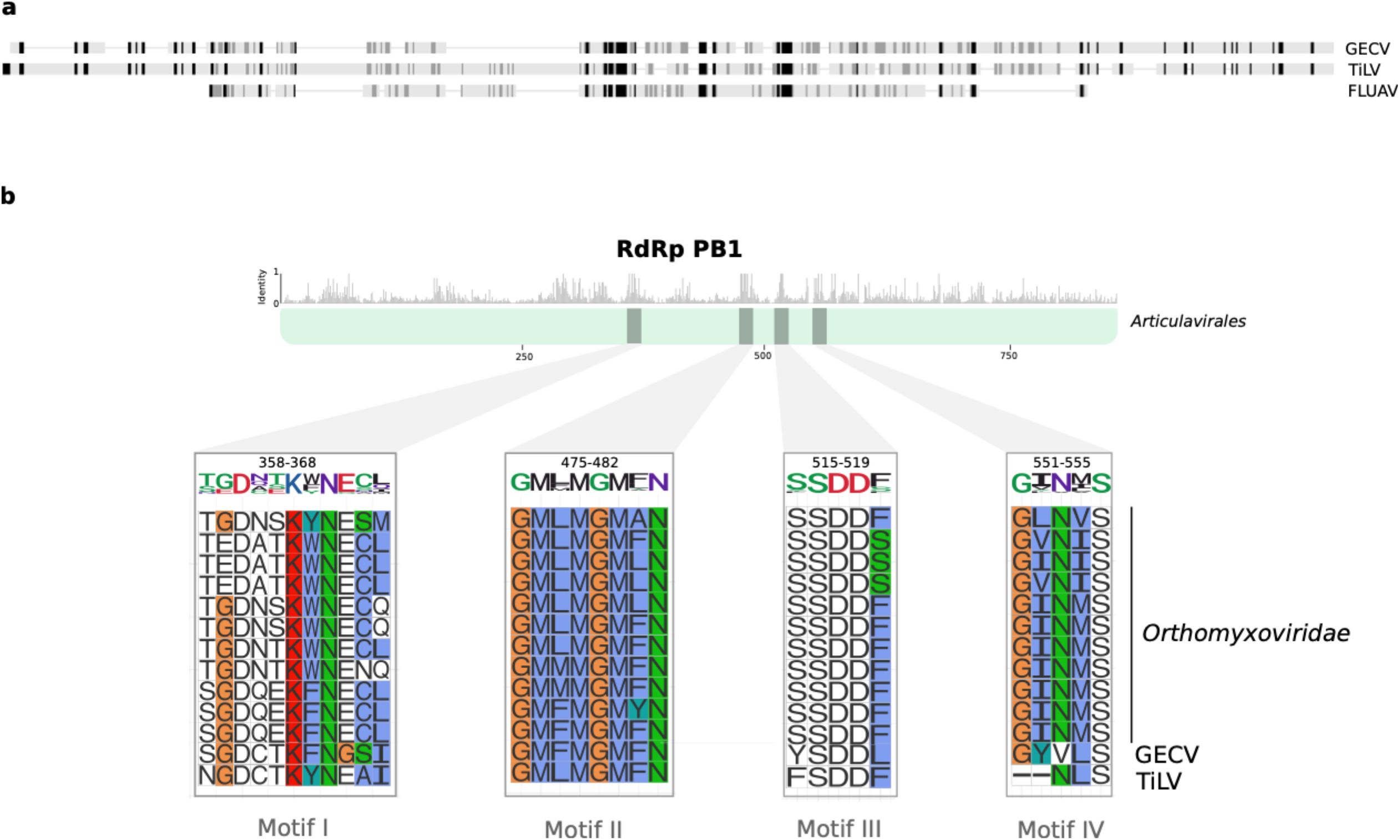
Conserved motifs in the RdRp subunit PB1 from the order *Articulavirales*. (**a**) Comparison of the GECV RdRp sequence with the full-length PB1 sequence of TiLV and FLUAV. (**b**) Top panel shows the mean pairwise identity over all pairs in the column across the multiple sequence alignment. The bottom panel depicts the individual motifs. The original amino acid residue position and standard logos are displayed in the top of each motif; the size of each character represents the level of sequence conservation. Amino acid residues in the alignment are coloured according to the Clustal colouring scheme.

### Host association and in vitro validation

GECV was initially identified in the pooled sequencing library comprising a mix of several Australian reptile species. To identify the exact host species, we screened each individual species sample separately using RT-PCR and Sanger sequencing. As a result, we detected the presence of the novel GECV RdRp sequence in liver tissue of *G. lauta* (paratype QM J96622) (Figure S1), a gecko species native to north-western Queensland and the north-eastern Northern territory in Australia [30].

## Discussion

Advances in protein modelling and sequence analysis based on structural comparisons with well-characterized protein templates constitute an attractive approach for the identification of highly divergent RNA viruses [27]. As viral proteins such as the RdRp play a central role on transcription and replication of RNA viruses, it is expected that structures and key motifs for catalytic functionality will be relatively well conserved throughout evolutionary history [31,32]. Based on this premise, it is expected that template-based protein structure modelling could be a powerful tool in the identification of highly divergent viruses [7,27,33]. Accordingly, we used protein structural similarity in combination with sequence and a profile similarity to identify a novel and divergent RNA virus in an Australian gecko (*G. lauta*).

We obtained a confident predicted 3D model for the RdRp of GECV based on its structural similarity with the RdRp subunit PB1 of influenza virus (family *Orthomyxoviridae*) (Figure 1a-b; Table 1). Although the structural data suggested that GECV belonged to the family *Orthomyxoviridae* (order *Articulavirales*) [27], additional sequence analysis revealed a closer relationship to members of the family *Amnoonviridae* (Figure 1c). In this context it is important to recall that biases in taxonomic assignment can occur because of the limited number of available proteins with known structures in the PDB. Although this is clearly a limitation, template-based approaches offer a tractable starting point for virus discovery and its taxonomic classification.

Although compromised by the large evolutionary distances involved, phylogenetic analysis among members of the order *Articulavirales* revealed that GECV was most closely related to TiLV, in turn suggesting that GECV is a novel and divergent genus within the *Amnoonviridae*. To date, the family *Amnoonviridae* has only been detected in fish [14], such that the discovery of GECV expands the host range of this family. Indeed, given the distance between the TiLV and GECV viruses, we can expect that further uncharacterised diversity exists in the family *Amnoonviridae* especially in fish and reptiles, and that more studies using the form of genomic surveillance performed here will reveal a far greater diversity of negative-sense RNA viruses [6,34].

Comparisons of the RdRp subunit PB1 from different articulaviruses revealed the presence of four well conserved motifs in GECV, broadly consistent with observations made for TiLV [14]. As suggested by several studies, motifs I-IV are critically implicated in the catalytic activity of PB1 [35,36]. Despite minor variations, we identified the SDD (serine-aspartic acid-aspartic acid) sequence in motif III that is presumed to be essential for protein functionality in FLUV [35,36]. Hence, the presence of well conserved motifs I-IV across the order *Articulavirales* may constitute effective molecular fingerprints for these viruses. Unfortunately, the marked lack of sequence similarity meant we did not recover any conclusive evidence regarding presence of other genome segments in GECV. Further studies that include sequencing, microscopy, and cell culture techniques, are therefore required to fully characterize the genome of this novel virus.

The identification of a novel virus in an Australian gecko (*G. lauta*) highlights the importance of virus surveillance in native species. Although GECV was detected in liver tissue, we currently cannot draw any conclusions regarding its pathogenic potential and impact on the health of *G. lauta*, particularly since a limited number of individuals were collected and all were apparently healthy. Additional research is therefore needed to establish the type of biological interaction between GECV and *G. lauta*. While a previous study reported the isolation of the arbovirus Charleville virus (family *Rhabdoviridae*) in *G. australis* (possibly *G. dubia* based on its distribution) collected in Queensland [36,37], this is the first report of a divergent articulavirus in reptiles. Taken together, these findings hint at a hidden diversity of RNA viruses in reptiles that remains to be characterized.

## Supplementary Materials.

**Figure S1.** PCR detection and host association of GECV. (a-b) Agarose gels electrophoresis showing PCR products from two sets of primers that target a region in the PB1 gene segment (RdRp). Samples correspond to (c) liver tissue from seven different reptile species. A 355 bp PCR product was only amplified in *G. lauta.*

**Table S1.** Summary of the contig alignment to genomic segments of TiLV using DIAMOND. The relative abundance of each transcript was also calculated (see Methods).

**Table S2.** List of virus sequences used in the phylogenetic analysis. All sequences correspond to the PB1 protein.

**Table S3.** Set of primers used for PCR and Sanger sequencing reactions.

## Author Contributions

Conceptualization, E.C.H.; methodology, A.S.O.-B., E.C.H., and J.-S.E.; formal analysis, A.S.O.-B.; investigation, A.S.O.-B., E.C.H., and J.-S.E.; resources, C.M., J.-S.E and E.C.H.; writing—original draft preparation A.S.O.-B.; writing—review and editing E.C.H., J.-S.E. and C.M.; visualization, A.S.O.-B.; supervision, E.C.H. All authors have read and agreed to the published version of the manuscript.

## Acknowledgments

None.

## References

1. Thermes, C. Ten years of next-generation sequencing technology. Trends Genet. 2014, 30, 418–426.

2. Shi, M.; Lin, X.-D.; Chen, X.; Tian, J.-H.; Chen, L.-J.; Li, K.; Wang, W.; Eden, J.-S.; Shen, J.-J.; Liu, L.; Holmes, E.C.; Zhang, Y.-Z. The evolutionary history of vertebrate RNA viruses. Nature 2018, 556, 197–202.

3. Zhang, Y.-Z.; Chen, Y.-M.; Wang, W.; Qin, X.-C.; Holmes, E.C. Expanding the RNA virosphere by unbiased metagenomics. Annu. Rev. Virol. 2019, 6, 119–139.

4. Zhang, Y.-Z.; Shi, M.; Holmes, E.C. Using metagenomics to characterize an expanding virosphere. Cell 2018, 172, 1168–1172.

5. Rose, R.; Constantinides, B.; Tapinos, A.; Robertson, D.L.; Prosperi, M. Challenges in the analysis of viral metagenomes. Virus Evol. 2016, 2, vew02,.

6. Shi, M.; Lin, X.-D.; Vasilakis, N.; Tian, J.-H.; Li, C.-X.; Chen, L.-J.; Eastwood, G.; Diao, X.-N.; Chen, M.-H.; Chen, X.; Qin, X.-C.; Widen, S.G.; Wood, T.G.; Tesh, R.B.; Xu, J.; Holmes, E.C.; Zhang, Y.-Z. Divergent viruses discovered in arthropods and vertebrates revise the evolutionary history of the *Flaviviridae* and related viruses. J. Virol. 2016, 90, 659–669.

7. Deng, H.; Jia, Y.; Zhang, Y. Protein structure prediction. Int. J. Mod. Phys. B 2018, 32.

8. Holmes, E.C. What does virus evolution tell us about virus origins? J. Virol. 2011, 85, 5247–5251.

9. Bamford, D.H.; Grimes, J.M.; Stuart, D.I. What does structure tell us about virus evolution? Curr. Opin. Struct. Biol. 2005, 15, 655–663.

10. Benson, S.D.; Bamford, J.K.H.; Bamford, D.H.; Burnett, R.M. Does common architecture reveal a viral lineage spanning all three domains of life? Mol. Cell 2004, 16, 673–685.

11. Rice, G.; Tang, L.; Stedman, K.; Roberto, F.; Spuhler, J.; Gillitzer, E.; Johnson, J.E.; Douglas, T.; Young, M. The structure of a thermophilic archaeal virus shows a double-stranded DNA viral capsid type that spans all domains of life. Proc. Natl. Acad. Sci. U.S.A. 2004, 101, 7716–7720.

12. Baker, D.; Sali, A. Protein structure prediction and structural genomics. Science 2001, 294, 93–96.

13. Shi, M.; Lin, X.D.; Tian, J.H.; Chen, L.J.; Chen, X.; Li, C.X.; Qin, X.C.; Li, J.; Cao, J.P.; Eden, J.S.; Buchmann, J.; Wang, W.; Xu, J.; Holmes, E.C.; Zhang, Y.Z. Redefining the invertebrate RNA virosphere. Nature 2016, 540, 539–543.

14. Bacharach, E.; Mishra, N.; Briese, T.; Zody, M.C.; Kembou Tsofack, J.E.; Zamostiano, R.; Berkowitz, A.; Ng, J.; Nitido, A.; Corvelo, A.; Toussaint, N.C.; Abel Nielsen, S.C.; Hornig, M.; Del Pozo, J.; Bloom, T.; Ferguson, H.; Eldar, A.; Lipkin, W.I. Characterization of a novel orthomyxo-like virus causing mass die-offs of Tilapia. mBio 2016, 7, e00431–16.

15. Jansen, M.D.; Dong, H.T.; Mohan, C.V. Tilapia Lake Virus: a threat to the global Tilapia industry? Rev. Aquac. 2019, 11, 725–739.

16. Pulido, L.L.H.; Mora, C.M.; Hung, A.L.; Dong, H.T.; Senapin, S. Tilapia Lake Virus (TiLV) from Peru is genetically close to the Israeli isolates. Aquaculture 2019, 510, 61–65.

17. Ahasan, M.S.; Keleher, W.; Giray, C.; Perry, B.; Surachetpong, W.; Nicholson, P.; Al-Hussinee, L.; Subramaniam, K.; Waltzek, T.B. Genomic characterization of Tilapia Lake Virus Iiolates recovered from moribund Nile Tilapia (*Oreochromis niloticus*) on a farm in the United States. Microbiol. Resour. Announc. 2020, 9, e01368–19.

18. Subramaniam, K.; Ferguson, H.W.; Kabuusu, R.; Waltzek, T.B. Genome sequence of Tilapia Lake Virus associated with syncytial hepatitis of Tilapia in an Ecuadorian aquaculture facility. Microbiol. Resour. Announc. 2019, 8, e00084–19.

19. Al-Hussinee, L.; Subramaniam, K.; Ahasan, M.S.; Keleher, B.; Waltzek, T.B. Complete genome sequence of a Tilapia Lake Virus isolate obtained from Nile tilapia (*Oreochromis Niloticus*). Genome Announc. 2018, 6, e00580–18.

20. Payne, S. Family *Orthomyxoviridae*. In Viruses; Elsevier, 2017; pp 197–208.

21. Bolger, A.M.; Lohse, M.; Usadel, B. Trimmomatic: a flexible trimmer for Illumina sequence data. Bioinformatics 2014, 30, 2114–2120.

22. Grabherr, M.G.; Haas, B.J.; Yassour, M.; Levin, J.Z.; Thompson, D.A.; Amit, I.; Adiconis, X.; Fan, L.; Raychowdhury, R.; Zeng, Q.; Chen, Z.; Mauceli, E.; Hacohen, N.; Gnirke, A.; Rhind, N.; di Palma, F.; Birren, B.W.; Nusbaum, C.; Lindblad-Toh, K.; Friedman, N.; Regev, A. full-length transcriptome assembly from RNA-Seq data without a reference genome. Nat. Biotechnol. 2011, 29, 644–652.

23. Li, B.; Dewey, C.N. RSEM: Accurate transcript quantification from RNA-Seq data with or without a reference genome. BMC Bioinformatics 2011, 12, 323.

24. Altschul, S.F.; Gish, W.; Miller, W.; Myers, E.W.; Lipman, D.J. Basic Local Alignment Search Tool. J. Mol. Biol. 1990, 215, 403–410.

25. Buchfink, B.; Xie, C.; Huson, D.H. Fast and sensitive protein alignment using DIAMOND. Nat. Methods 2015, 12, 59–60.

26. Rice, P.; Longden, L.; Bleasby, A. EMBOSS: The European Molecular Biology open software suite. Trends Genet. 2000, 16, 276–277.

27. Kelley, L.A.; Mezulis, S.; Yates, C.M.; Wass, M.N.; Sternberg, M.J.E. The Phyre2 web portal for protein modeling, prediction and analysis. Nat. Protoc. 2015, 10, 845–858.

28. Katoh, K.; Standley, D.M. MAFFT Multiple Sequence Alignment Software Version 7: improvements in performance and usability. Mol. Biol. Evol. 2013, 30, 772–780.

29. Nguyen, L.-T.; Schmidt, H.A.; von Haeseler, A.; Minh, B.Q. IQ-TREE: A fast and effective stochastic algorithm for estimating maximum-likelihood phylogenies. Mol. Biol. Evol. 2015, 32, 268–274.

30. Oliver, P.M.; Prasetya, A.M.; Tedeschi, L.G.; Fenker, J.; Ellis, R.J.; Doughty, P.; Moritz, C. Crypsis and convergence: integrative taxonomic revision of the *Gehyra Australis* group (Squamata: Gekkonidae) from Northern Australia. PeerJ 2020, 2020, e7971.

31. Zanotto, P.M. de A.; Gibbs, M.J.; Gould, E.A.; Holmes, E.C. A reevaluation of the higher taxonomy of viruses based on RNA polymerases. J. Virol. 1996, 70, 6083–6096..

32. Ng, K.K.S.; Arnold, J.J.; Cameron, C.E. Structure-function relationships among RNA-dependent RNA polymerases. Curr. Top. Microbiol. Immunol. 2008, 320, 137–156.

33. Fiser, A. Template-based protein structure modeling. Methods in molecular biology (Clifton, N.J.). Humana Press, Totowa, NJ 2010, pp 73–94.

34. Li, C.-X.; Shi, M.; Tian, J.-H.; Lin, X.-D.; Kang, Y.-J.; Chen, L.-J.; Qin, X.-C.; Xu, J.; Holmes, E.C.; Zhang, Y.-Z. Unprecedented genomic diversity of RNA viruses in arthropods reveals the ancestry of negative-sense RNA viruses. eLife 2015, 4, e05378.

35. Biswas, S.K.; Nayak, D.P. Mutational analysis of the conserved motifs of influenza A virus polymerase basic protein 1. J. Virol. 1994, 68, 1819–1826.

36. Chu, C.; Fan, S.; Li, C.; Macken, C.; Kim, J.H.; Hatta, M.; Neumann, G.; Kawaoka, Y. Functional analysis of conserved motifs in influenza virus PB1 protein. PLoS One 2012, 7, e36113.

