## Supplementary Material for "A divergent *Articulavirus* in an Australian gecko identified using meta-transcriptomics and protein structure comparisons"

**Supplementary Materials**

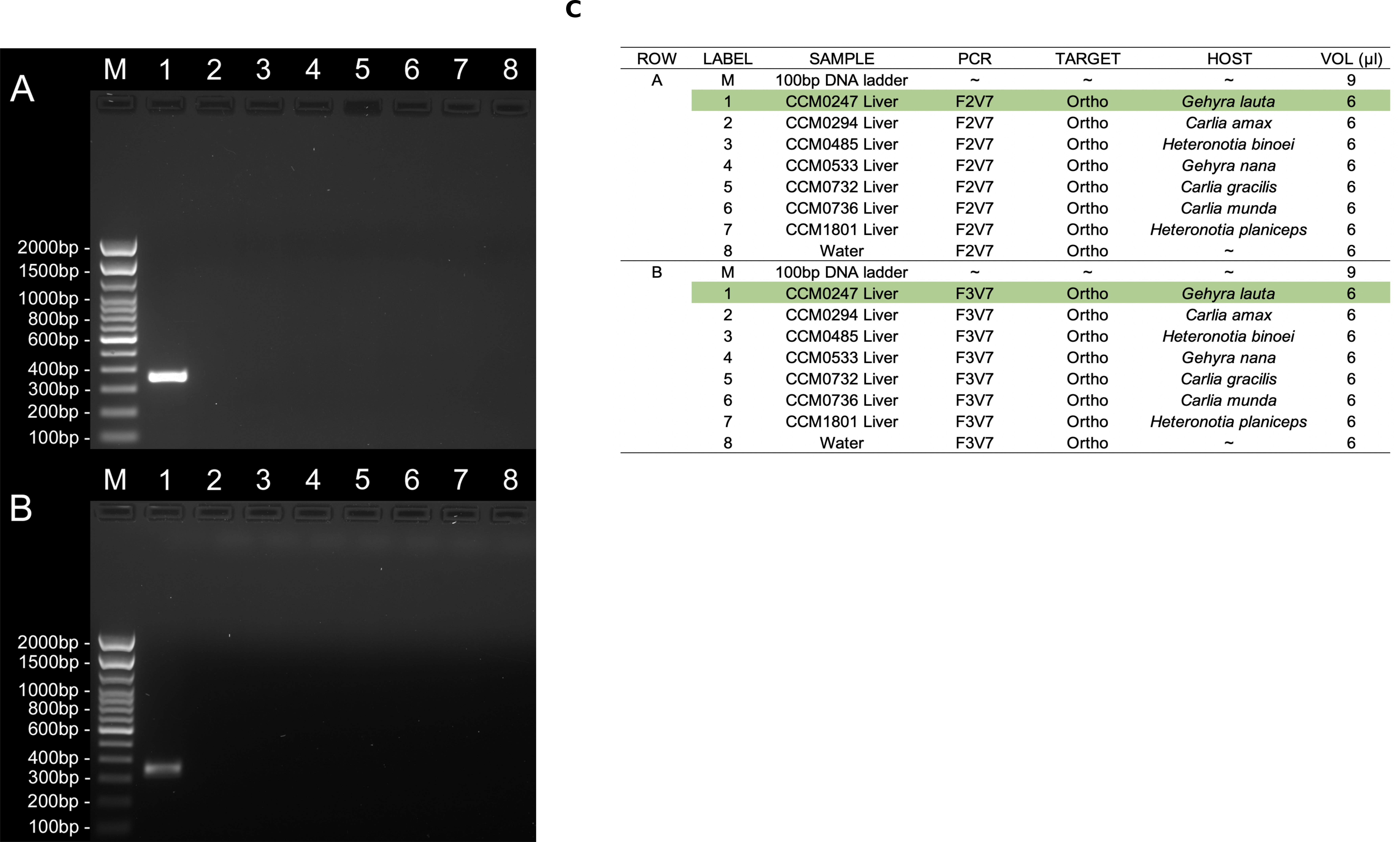

**Figure S1.** PCR detection and host association of GECV. (**a-b**) Agarose gels electrophoresis showing PCR products from two sets of primers that target a region in the PB1 gene segment (RdRp). Samples correspond to (**c**) liver tissue from seven different reptile species. A 355 bp PCR product was only amplified in *G. lauta.*

**Table S1.** Summary of the contig alignment to genomic segments of TiLV using DIAMOND. The relative abundance of each transcript was also calculated (see Methods).

**Table S2.** List of virus sequences used in the phylogenetic analysis. All sequences correspond to the PB1 protein.

| Species name | Acronym | GenBank access code | Family |
| --- | --- | --- | --- |
| *Blueberry mosaic associated virus* | BIMaV | YP_009449565 | *Ophioviridae* - Outgroup |
| *Montano orthohantavirus* | MTNV | YP_009361849 | *Hantaviridae* - Outgroup |
| *Bayou orthohantavirus* | BAYV | YP_009505596 | *Hantaviridae* - Outgroup |
| *Influenza A virus* | FLUAV | YP_009118628 | *Orthomyxoviridae* |
| *Influenza B virus* | FLUBV | NP_056657 | *Orthomyxoviridae* |
| *Influenza D virus* | FLUDV | YP_009449556 | *Orthomyxoviridae* |
| *Influenza C virus* | FLUCV | YP_089653 | *Orthomyxoviridae* |
| *Salmon isavirus* | ISAV | YP_145804 | *Orthomyxoviridae* |
| *Quaranfil quaranjavirus* | QRFV | YP_009508043 | *Orthomyxoviridae* |
| *Thogoto thogotovirus* | THOV | YP_145794 | *Orthomyxoviridae* |
| *Tilapia tilapinevirus* | TiLV | YP_009246481 | *Amnoonviridae* |
| *Dhori thogotovirus* | DHOV | YP_009352882 | *Orthomyxoviridae* |
| *Oz virus* | OZV | YP_009553280 | *Orthomyxoviridae* |
| *Wellfleet Bay virus* | WFBV | YP_009110686 | *Orthomyxoviridae* |
| *Johnston Atoll quaranjavirus* | JAV | YP_009665204 | *Orthomyxoviridae* |

**Table S3.** Set of primers used for PCR and Sanger sequencing reactions.

| Primer | Nucleotide sequence (5’-3’) | Tm (ºC) | Anneal (ºC) | Amplicon size (bp) |
| --- | --- | --- | --- | --- |
| F2V7_136F | ACTGCACCAAGTTCAACGGA | 61.8 | 65.2 | 355 |
| F2V7_490R | GTGAGTGGGTCCATCTTGCA | 61.7 |  |  |
| F3V7_396F | TGCAGTAGAGCGGAGGTAGA | 61.6 | 64.4 | 355 |
| F3V7_711R | ATCCGAGCCCAGCATATCTC | 60.9 |  |  |
