## Supplementary Figure 1 for "A divergent *Articulavirus* in an Australian gecko identified using meta-transcriptomics and protein structure comparisons"

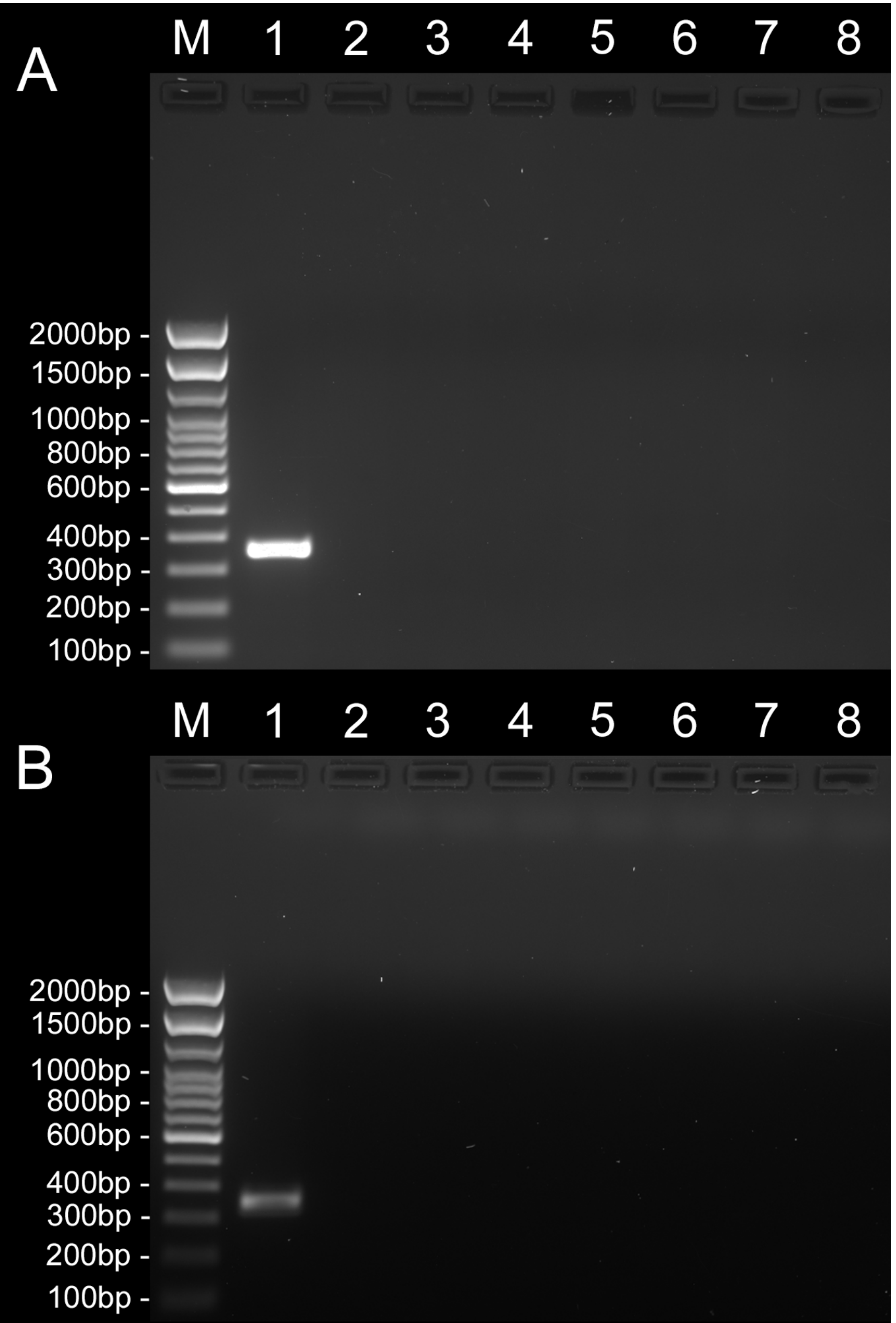


**C**

| ROW | LABEL | SAMPLE | PCR | TARGET | HOST | VOL (µl) |
|---|---|---|---|---|---|---|
| A | M | 100bp DNA ladder | ~ | ~ | ~ | 9 |
| | 1 | CCM0247 Liver | F2V7 | Ortho | *Gehyra lauta* | 6 |
| | 2 | CCM0294 Liver | F2V7 | Ortho | *Carlia amax* | 6 |
| | 3 | CCM0485 Liver | F2V7 | Ortho | *Heteronotia binoei* | 6 |
| | 4 | CCM0533 Liver | F2V7 | Ortho | *Gehyra nana* | 6 |
| | 5 | CCM0732 Liver | F2V7 | Ortho | *Carlia gracilis* | 6 |
| | 6 | CCM0736 Liver | F2V7 | Ortho | *Carlia munda* | 6 |
| | 7 | CCM1801 Liver | F2V7 | Ortho | *Heteronotia planiceps* | 6 |
| | 8 | Water | F2V7 | Ortho | ~ | 6 |
| B | M | 100bp DNA ladder | ~ | ~ | ~ | 9 |
| | 1 | CCM0247 Liver | F3V7 | Ortho | *Gehyra lauta* | 6 |
| | 2 | CCM0294 Liver | F3V7 | Ortho | *Carlia amax* | 6 |
| | 3 | CCM0485 Liver | F3V7 | Ortho | *Heteronotia binoei* | 6 |
| | 4 | CCM0533 Liver | F3V7 | Ortho | *Gehyra nana* | 6 |
| | 5 | CCM0732 Liver | F3V7 | Ortho | *Carlia gracilis* | 6 |
| | 6 | CCM0736 Liver | F3V7 | Ortho | *Carlia munda* | 6 |
| | 7 | CCM1801 Liver | F3V7 | Ortho | *Heteronotia planiceps* | 6 |
| | 8 | Water | F3V7 | Ortho | ~ | 6 |
